# Characterization of Natural Killer Cell-Associated Prognostic Signature Reveals Immune Landscape and Prognostic Value in Uterine Corpus Endometrial Carcinoma (UCEC)

**DOI:** 10.64898/2026.11.24.768788

**Authors:** Qin Liu, Xiaoli Chen, Kaibo Zhu, Minghua Yu, Qin Liu

**Affiliations:** Department of Pathology, Zhejiang University School of Medicine Women’s Hospital, Hangzhou, Zhejiang Province, China, 310006

**Keywords:** Natural killer cell, UCEC, tumor microenvironment, prognosis, immunotherapy

## Abstract

**Background:** Natural killer (NK) cells are critical in eliminating virus-infected and cancer cells. Uterine corpus endometrial carcinoma (UCEC) is a common malignancy in women, and the role of NK cells in UCEC has attracted significant attention. This study aims to develop an NK cell-related gene signature (NRGs) to improve immunotherapy response and prognosis evaluation in UCEC.

**Methods:** Consensus clustering classified UCEC samples based on NRGs. Univariate Cox regression, differential expression, and Lasso regression were used to build a prognostic model. Differences in tumor microenvironment (TME), immune checkpoints, and clinical features were compared between high- and low-risk groups.

**Results:** High NK cell infiltration was associated with a favorable prognosis. Eleven key NRGs were identified, and consensus clustering divided patients into two clusters: Cluster 1 (C1) had better prognosis and higher immune cell infiltration. A nine-NRG signature (CEACAM1, IFNG, PTPN6, KNSTRN, XCL1, IL2RG, RAC3, MICB, CCL3) was developed. The high-risk group had more advanced-stage cases, higher-grade tumors, and serous carcinoma compared to the low-risk group (P < 0.05).

**Conclusions:** The 9-NRGs signature demonstrates strong predictive performance and offers valuable insights for future immuno-oncology research.

## Introduction

Endometrial cancer is one of the most common malignancies in the female reproductive system, with over 320,000 new cases of UCEC reported globally each year [1]. I Several factors, including socioeconomic progress, dietary changes, increasing obesity and diabetes rates, and population aging, have contributed to the rising incidence of UCEC [2, 3]. While early-stage UCEC generally has a favorable prognosis, advanced stages pose significant risks to women’s health [4]. Conventional treatments, such as surgery, chemotherapy, radiotherapy, and hormonal therapy, often fail to achieve complete disease eradication. Immunotherapy, particularly immune checkpoint inhibitors in dMMR or MSI-H tumors, has shown promising results[5, 6]. Combination strategies and the identification of predictive biomarkers offer potential for further improving treatment outcomes.

NK cells comprise approximately 15% of circulating lymphocytes and play a crucial role in the innate immune response through early cytokine and chemokine release and direct elimination of target cells without prior sensitization[7]. They offer distinct advantages over T cells in combating solid tumors, including enhancing responses to radioimmunotherapy and chemotherapy [8, 9]. Despite lower levels of tumor immune infiltration[10], NK cells hold significant prognostic value [11-13]. Recent single-cell sequencing analyses have identified NK cells as a major component of tumor-infiltrating lymphocytes (TILs) in UCEC patients [14, 15]. With advances in immunotherapy, combination strategies targeting NK cells alongside conventional treatments show promise for improving clinical outcomes[16]. Thus, understanding the role of NRGs in UCEC, particularly in relation to diagnosis and prognosis, is essential for developing effective immunotherapeutic approaches.

In the study, we conducted a comprehensive analysis of TCGA-UCEC cohort to identify 9 robust NRGs for constructing a prognostic model. Using these NRGs, we developed a prognostic model to evaluate their association with the immune microenvironment and potential implications for immunotherapy. Our study aims to highlight the significance of NRGs in assessing the prognosis of UCEC patients by integrating genomic data, thus supporting the development of novel tools that can enhance treatment options.

## Methods

### The acquisition of NK cells-related genes (NRGs) and data sources

A total of 244 NRGs that are associated with NK cells were obtained from the immport portal website(https://www.immport.org/resource) and the Molecular Signature Database (MSigDB). We obtained UCEC datasets and clinical information from the TCGA database (https://portal.gdc.cancer.gov/). The expression profiles, initially in FPKM format, were converted to TPM format [17]. Gene expression data were then normalized using a log2 transformation (log2(exp + 1)).

### Evaluation of the immune cell infiltration status in UCEC

CIBERSORT is a tool for immune infiltration analysis that assesses the abundance of 22 immune cell types in tumor and normal tissues. In this study, we used the “e1701” R package to perform CIBERSORT analysis, iterating 1000 times to calculate immune infiltration levels in UCEC.

### Identification of subtypes based on NRGs

We performed univariate Cox analysis using the coxph function to identify prognostic NRGs. Expression data of 11 prognostic NRGs were then used for non-negative matrix factorization (NMF) analysis. To explore signaling mechanisms between the two NRG-based subtypes, we conducted Gene Set Enrichment Analysis (GSEA) (version 4.0; https://www.gsea-msigdb.org/, accessed 26 October 2022). Enriched biological functions were identified using C2 gene sets (c2.cp.kegg.v7.1.symbols.gmt) from the Molecular Signatures Database (MSigDB). Statistical significance was set at P < 0.05 and FDR < 0.25, with 1000 permutations per analysis.

### Analysis of Somatic Mutation

The somatic mutation data for the UCEC samples were obtained from the TCGA GDC Data Portal. To visualize the differences in mutated genes in the two subtypes based on NRGs, we generated waterfall plots using the “Maftools” package in the R software.

### Construction of the novel prognostic model based on NRGs

The LASSO regression algorithm was implemented using the “glmnet” R package. Finally, 11 NRGs that exhibited significant correlations with overall survival (OS) in UCEC patients were selected to construct a novel prognostic model. The risk score was calculated using the formula: Riskscore=(-0.1740)*CEACAM1+(-0.0280)*IFNG+(-0.1647)*PTPN6+(0.0878)*KNSTRN+(-0.0378)*XCL1+(-0.0674)*IL2RG+(0.0637)*RAC3+(0.1611)*MICB+(-0.0916)*CCL3. Based on the median risk score, all subjects were classified into high and low risk groups. Kaplan-Meier plotter analysis was utilized to compare OS in high and low risk groups. Time-dependent receiver operating characteristic (ROC) curve analysis was performed to assess the predictive accuracy of the prognostic model for overall survival (OS) at various time points. Furthermore, univariate and multivariate Cox regression analyses were conducted, considering common factors associated with UCEC.

## Results

### Identification of prognostic NRGs in UCEC

We first applied the CIBERSORT algorithm to assess the infiltration levels of 22 immune cell types using TCGA-UCEC data. Kaplan-Meier analysis indicated that high NK cell infiltration correlated with a favorable prognosis (Fig. 1A). The expression patterns of 36 differentially expressed NRGs were visualized using heatmaps and volcano plots (Fig. 1B-D). Protein-protein interactions among these NRGs were explored via the STRING database (Fig. 1E). Subsequently, univariate Cox regression analysis identified 18 prognostic NRGs from the 36 differentially expressed candidates (Fig. 1F).

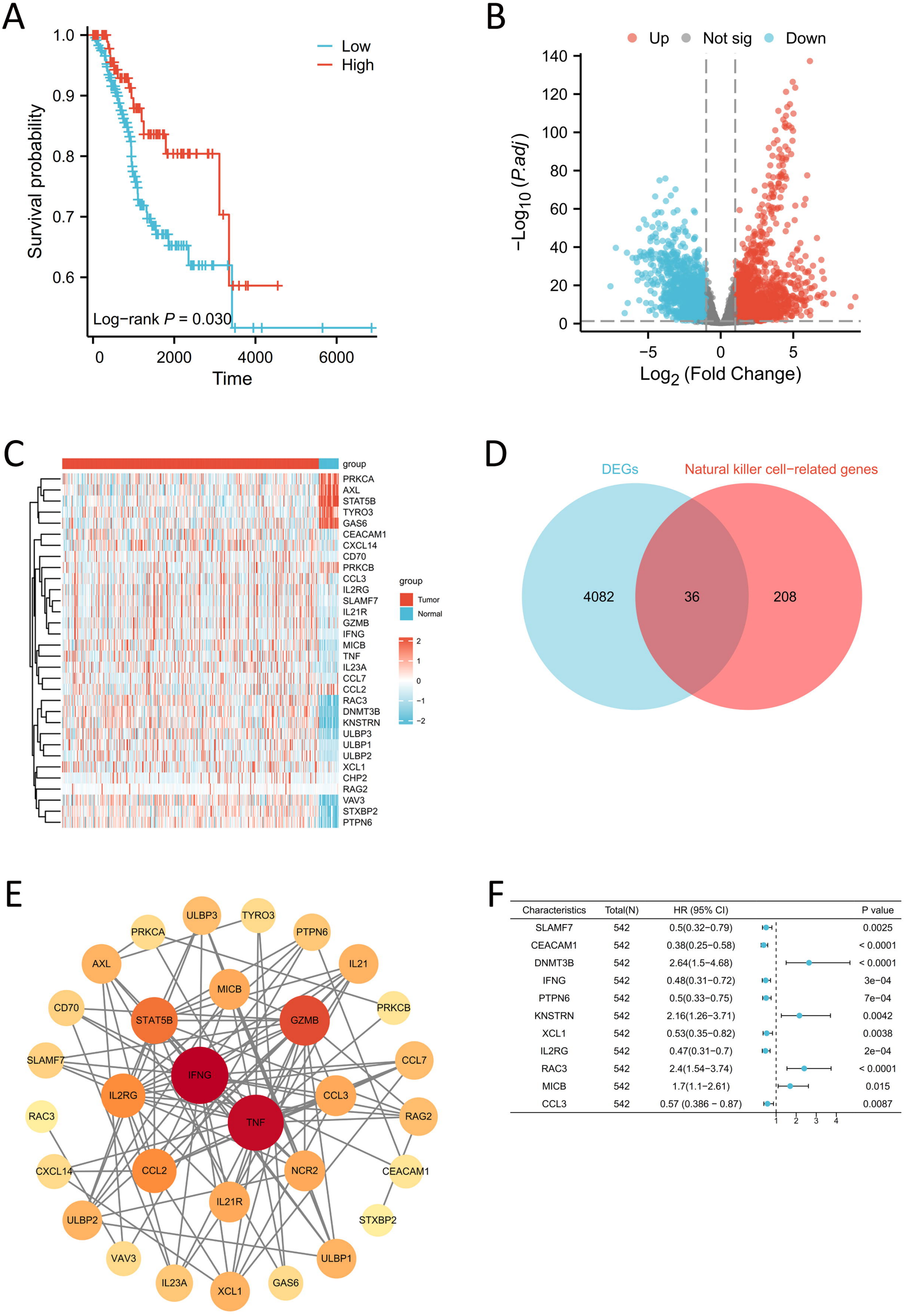

### Consensus Clustering based on 18 prognostic NRGs and subsequent pathway analysis

Using the 18 prognostic NRGs, the RConsensusClusterPlus package identified two distinct clusters in the TCGA-UCEC dataset: Cluster 1 (188 cases) and Cluster 2 (175 cases) (Fig. 2A-D). Cluster 2 was associated with significantly worse prognosis compared to Cluster 1 (log-rank p < 0.0001, Fig. 2E). GSEA revealed that immune-related pathways were primarily enriched in Cluster 1 (Fig. 2F). KEGG analysis of differentially expressed genes (DEGs) between the clusters showed enrichment in pathways such as Cytokine-cytokine receptor interaction, Chemokine signaling, NK cell-mediated cytotoxicity, and Primary immunodeficiency (Fig. 2G-H).

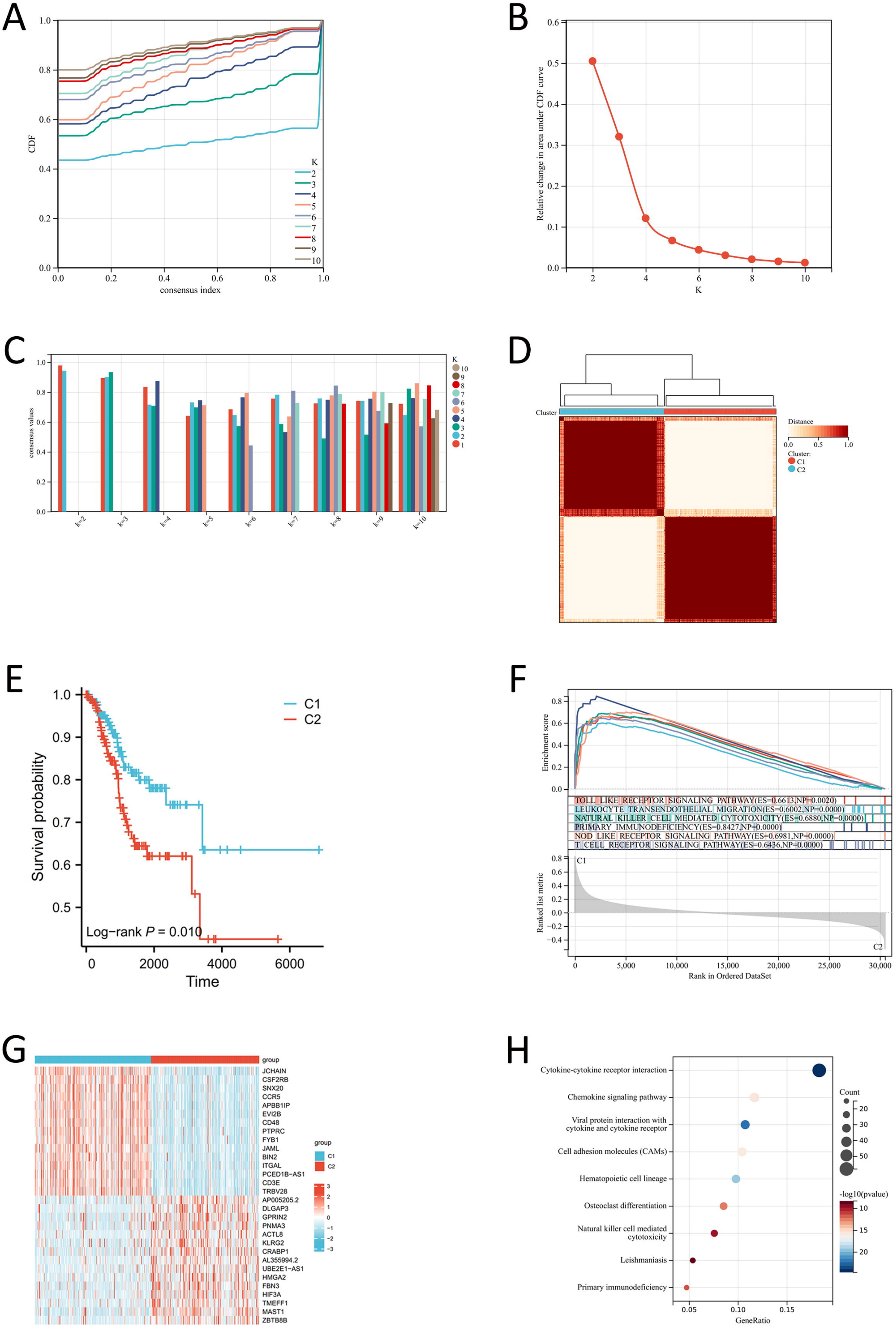

### Comparison of Somatic Mutations and Tumor Immune Microenvironment in two clusters

Infiltration of plasma cells, CD8 T cells, activated memory CD4 T cells, and regulatory T cells was significantly higher in Cluster 1 than in Cluster 2 (Fig. 3A-B). Somatic mutation analysis showed a higher frequency of PTEN mutations in Cluster 1, whereas P53 mutations were more frequent in Cluster 2 (Fig. 3C-D). Immune chemokines such as CCL5, CXCL9, IL1B, TNFSF4, IFNG, HMGB1, TLR4, TNFRSF14, TNFRSF4, CCR2, CCR5, CXCR3, ICAM1, and ITGB were upregulated in Cluster 1 (Fig. 3E). To assess the impact of NRGs on the immune microenvironment, we compared the tumor microenvironment between the clusters. Cluster 1 exhibited higher stromal, immune, and ESTIMATE scores than Cluster 2 (Fig. 3F), suggesting an immune-hot phenotype in Cluster 1 and an immune-cold phenotype in Cluster 2.

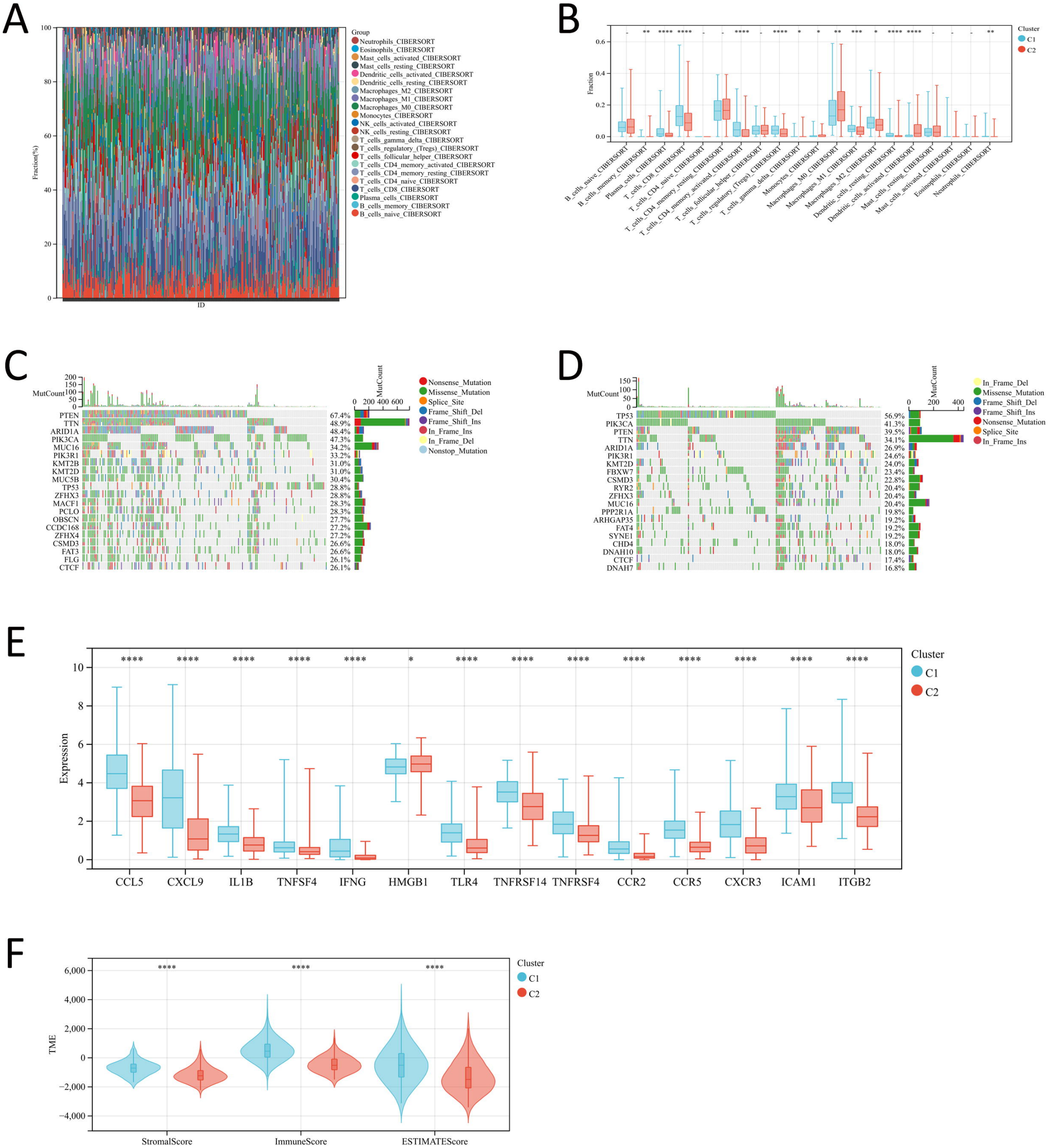

### Construction of Prognostic model based on NRGs

A prognostic model was developed using LASSO analysis, resulting in a 9-NRG signature (Fig. 4A-C). Patients were stratified into high- and low-risk subgroups based on an optimal cut-off for risk scores. Kaplan-Meier analysis indicated poorer prognosis in the high-risk subgroup (Fig. 4D). ROC curve analysis revealed AUC values of 0.674, 0.699, and 0.756, demonstrating the predictive value of the risk scores (Fig. 4E). Multivariate Cox regression analysis confirmed that the risk score from the TCGA-UCEC dataset is an independent prognostic factor (Fig. 4F).

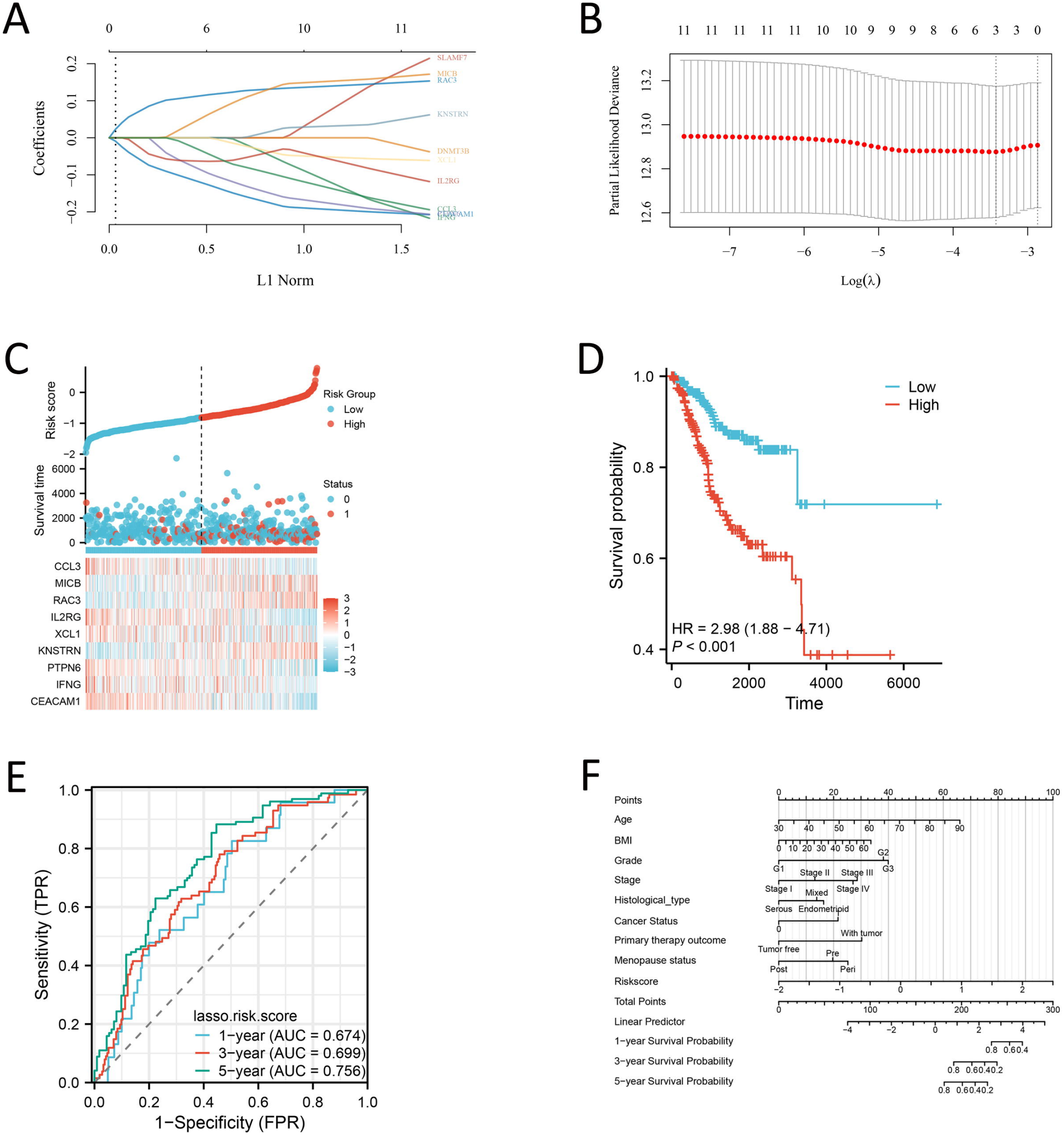

### High-and low-risk groups demonstrate distinctive patterns of immune cell infiltration and immune gene expression

Gene Set Enrichment Analysis (GSEA) revealed prominent enrichment of immune-related pathways in the low-risk group (Fig. 5A). Comparison of HLA family gene and immune checkpoint marker expression between high- and low-risk groups showed significantly higher expression of antigen-presentation and immune checkpoint genes in the low-risk group (Fig. 5B-D). The high-risk group was associated with a higher proportion of advanced-stage, high-grade tumors, and serous carcinoma (P < 0.05) (Fig. 5E).

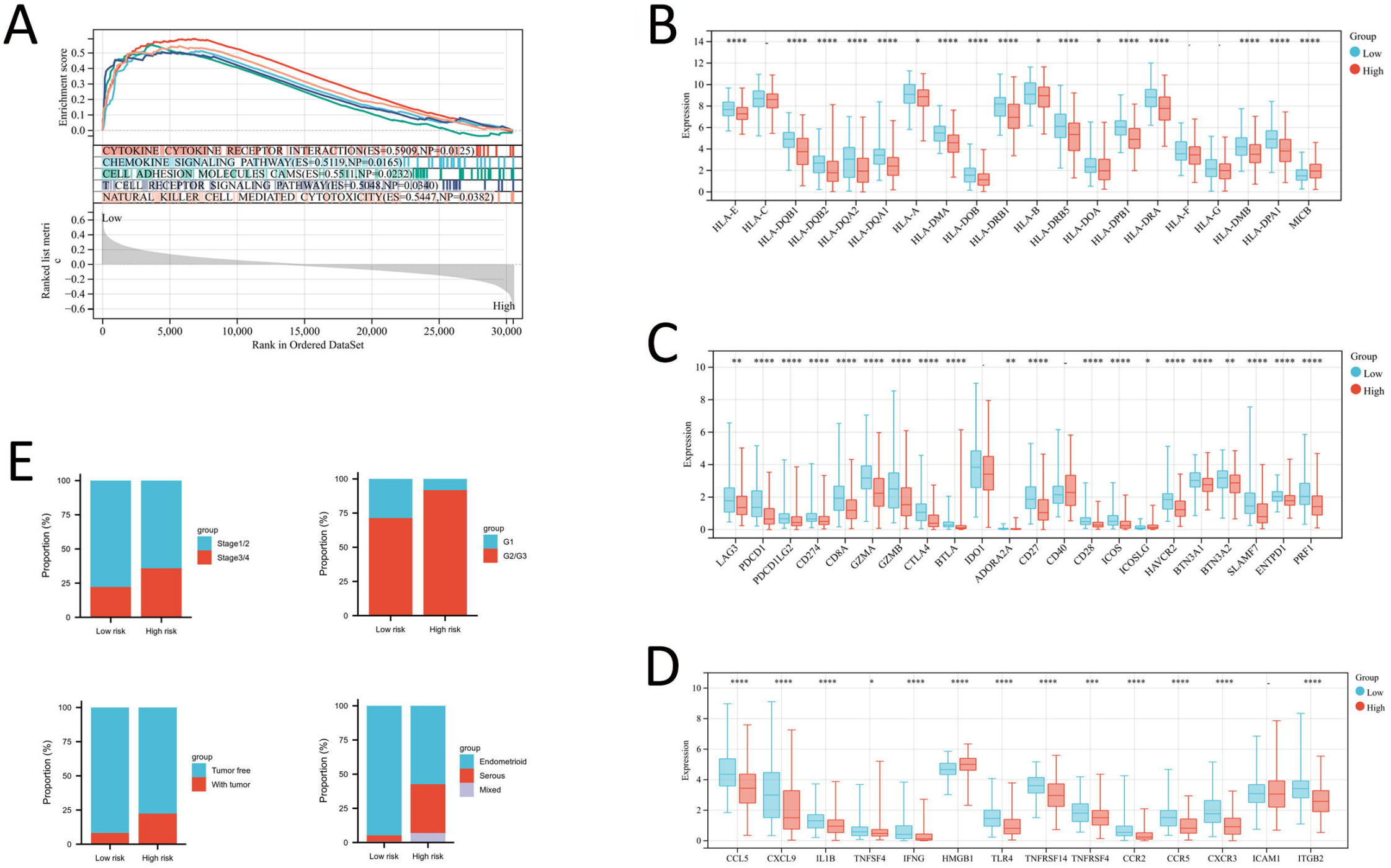

## Discussion

UCEC is one of the most common gynecologic malignancies globally, with rising incidence in recent years. Traditionally, endometrial cancer was classified into two types: type I (estrogen-dependent) and type II (estrogen-independent) [17, 18]. However, molecular profiling has led to a more refined classification, with The Cancer Genome Atlas (TCGA) identifying four distinct subtypes: POLE-mutated, microsatellite instability hypermutated, copy-number low, and copy-number high [19-21]. These subtypes differ in clinical behavior and treatment response, informing personalized therapy decisions. Recent advances, including the integration of immunotherapies, have further shaped UCEC management[22, 23]. Despite this progress, prognosis evaluation based on histological subtypes, molecular classification, or staging remains limited, underscoring the need for more accurate prognostic models to predict treatment outcomes.

Natural Killer (NK) cells are crucial components of the innate immune system, playing a key role in tumor surveillance and growth inhibition [24]. Studies have shown that NK cells can directly lyse specific tumor cells [25, 26], and their depletion is associated with increased tumor incidence [27, 28]. Additionally, NK cells are involved in recruiting T cells to the tumor microenvironment, enhancing their functionality[29]. Elevated NK cell levels in the tumor microenvironment have been linked to favorable prognosis in various cancers, including gastric cancer, melanoma, triple-negative breast cancer, head and neck squamous cell carcinoma, colorectal carcinoma, and squamous cell lung cancer[30-34]. Given their role in modulating anti-tumor immunity, research is increasingly focusing on NK cell-related genes[35]. However, the molecular mechanisms underlying NK cell-mediated anti-tumor activity remain poorly understood, and no model based on NK cell-related genes has yet been established to predict prognosis or assess the immune microenvironment in UCEC.

In this study, we developed a signature of nine NRGs (CEACAM1, IFNG, PTPN6, KNSTRN, XCL1, IL2RG, RAC3, MICB, CCL3) to predict survival in UCEC patients. CEACAM1, a heterophilic ligand for TIM-3, plays a key role in T-cell inhibition and immune regulation, particularly in autoimmunity and anti-tumor immunity, including myeloid-derived suppressor cell-induced CD8+ T cell exhaustion[36, 37]. Its expression is often downregulated in UCEC, though its role in this context remains underexplored [38]. However, there is limited research on the function of CEACAM1 in UCEC. IFNG is a pivotal mediator of antitumor immunity, enhancing macrophage activation and the antiviral and antitumor effects of type I interferons. It also contributes to the efficacy of anti-CTLA4 therapy by affecting vessel perfusion and angiostatic activity [39, 40]. Further research is needed to clarify IFNG’s role in UCEC immunotherapy. As for PTPN6, a cytoplasmic phosphatase, regulates autoimmune diseases and inflammatory responses, including those involving IL-1R [41]. Its dysfunction is implicated in various inflammatory disorders, such as neutrophilic dermatosis [42]. KNSTRN, a mitotic spindle protein, has been linked to tumorigenesis and chemotherapy resistance, particularly in bladder and breast cancers, where it influences cell cycle progression and immune infiltration [43, 44]. XCL1, a C-class chemokine commonly referred to as lymphotactin, is secreted by T, NK, and NKT cells in response to infection and inflammation[45]. XCL1 can promote infiltration and activation of cytotoxic T cells, thereby exerting anti-tumor effects[46]. IL2RG plays a critical role in JAK-STAT signaling, boosting cellular immune responses and serving as a potential biomarker for assessing treatment efficacy in cancer [47-50] As a member of Rho GTPases family, RAC3 probably acts as an oncogene in various types of cancer. RAC3 is overexpressed in lung cancer, breast cancer, or bladder cancer[51-53]. Further studies have demonstrated that RAC3 mediated autophagy in bladder cancer cells through the PI3K/AKT/mTOR pathway[54]. MICB, induced by cellular stress such as genomic damage, is associated with metastasis and advanced tumor stages, although its prognostic significance remains insufficiently validated[55, 56].

Approximately 70% of UCEC cases are diagnosed at an early stage with a relatively favorable prognosis. However, some patients may miss the opportunity for early diagnosis and treatment due to neglecting early symptoms such as irregular vaginal bleeding and discharge. Recently, targeted therapies and immunotherapies have shown promising efficacy in UCEC treatment. For example, Yongxu Mu et al. reported that umbilical cord blood-derived NK cell therapy is highly effective for UCEC[57]. Thus, incorporating both the tumor microenvironment (TME) and NRGs is critical for developing an “ideal” prognostic model for UCEC. Our study demonstrates that NK cell levels within the TME are associated with improved patient prognosis. Previous studies have also identified NK cells as key components of tumor-infiltrating lymphocytes (TILs) in UCEC patients[14, 15]. These findings inspired us to develop a prognostic model based on NK cell-related genes in UCEC.

Immune checkpoints are key components of the tumor microenvironment (TME). Elevated expression in the low-risk group suggests that cancer cells in these patients rely on the PD-1/PD-L1 signaling pathway to evade immune surveillance. Our study found higher expression levels of PD-1, TIM3, CTLA4, and LAG3 in the low-risk group. Notably, these checkpoint receptors are also expressed on NK cells, indicating that NK cells could enhance the therapeutic efficacy of anti-PD-L1 monoclonal antibodies and boost the anti-tumor response in PD-L1-high tumors. The low-risk group exhibited an immune-hot phenotype, while the high-risk group showed an immune-cold phenotype. Thus, elevated immune checkpoint expression may correlate with better responsiveness to targeted therapies.

This study is the first to develop a prognostic model based on an NK-cell-related gene signature, which includes a limited number of genetic components and achieves a 5-year AUC nearing 0.8. Furthermore, this signature shows promise in predicting the response to immunotherapy in UCEC. However, several limitations must be addressed. First, the study is primarily based on bioinformatics analyses and requires further experimental and clinical validation. Additionally, the relatively small sample size of UCEC cases may introduce potential bias into the findings.

